# Figure–ground relationship of voices in musical structure modulates reciprocal frontotemporal connectivity

**DOI:** 10.64898/2025.12.22.696104

**Authors:** Chan Hee Kim, Jeong-Eun Seo, Jaeho Seol, Chun Kee Chung

## Abstract

When listening to polyphonic music, we often perceive a melody as the figure against the ground of accompanying sounds. However, with repeated exposure, this figure–ground relationship may naturally shift, allowing the melody to recede into the ground. In a previous study, we found the consistent pattern of frontotemporal connectivity for the “Twinkle, Twinkle, Little Star” (TTLS) melody in the headings of two variations (II and IV) in Mozart’s 12 Variations, K. 265, indicating that the TTLS melody, but not the different lower voices, was the figure. However, the frontotemporal connectivity pattern may change in the same phrases repeating in the two variations. In the current study, we examined how frontotemporal connectivity changes in the repeated phrases. In the results, the frontotemporal connectivity pattern between the two variations changed in the final phrase after repeated passages. This suggests that the shift in the figure–ground relationship persists, with the TTLS melody becoming less prominent while the lower voices become relatively more prominent. Additionally, frontotemporal connectivity was strongly correlated with temporofrontal connectivity in the opposite direction. Finally, our data indicate that TTLS melody-based and sensory-based processes in response to a switched figure–ground relationship, are incorporated into the bidirectional connections between frontotemporal and temporofrontal connectivity. Our study highlights the brain’s ability to reconfigure figure– ground relationships in the processing of musical voices.

## Introduction

Humans can identify a specific melody within homophonic and polyphonic music because it often appears in a higher pitch range, making it easily distinguishable (Fujioka et al., 2005; Trainor et al., 2014). This phenomenon aligns with the figure–ground concept in Gestalt psychology (Köhler, 1967; Wagemans et al., 2012). Similar to a visual stimulus (Supplementary Figure 1), the melody can serve as the “figure,” while other voices constitute the “ground.” Additionally, listeners can sometimes shift their attention during a phrase, perceiving the melody as the background while other voices being dominant instead (Ragert et al., 2014; Deutch, 2019). However, even when the figure–ground relationship favors the figure as the more perceptually dominant voice, the ground can remain perceptible (Bigand et al., 2000), and vice versa.

Musical structure, including pitch, tonality, and harmony, is learned through experience, understanding of musical structure facilitates the recognition and anticipation of patterns in familiar pieces (Narmour, 2000; Tillmann et al., 2000). When a familiar melody appears in a musical piece, it is easily recognized the figure in the musical structure. However, repeated exposure to the melody may alter the figure–ground relationship between the upper and lower voices (Taher et al., 2016), and this change may eventually lead to the natural collapse of the figure–ground relationship centered on the upper voice of the familiar melody.

In our previous study using Mozart’s 12 Variations, K. 265 (Kim et al., 2020), we observed that only frontotemporal connectivity between the left Heschl’s gyrus (HG) and left inferior frontal gyrus (IFG) changed in response to the presence or absence of the “Twinkle, Twinkle, Little Star” (TTLS) melody of “C5-C5-G5-G5-A5”. This connectivity pattern for the TTLS was observed across a target phrase (T) of 2.1 seconds at the beginning of each variation. However, if the figure–ground relationship shifts after repetitions, the connectivity strength can become inconsistent (Supplementary Figure 1C). The present study examined how frontotemporal connectivity for *Variations II* and *IV* (Figure 1) changes across four target phrases (T1–T4) featuring the TTLS melody. We hypothesized that: 1) If the connectivity pattern does not differ significantly between *Variations II* and *IV*, the TTLS melody remains the figure, with the lower voice serving as the ground; and 2) If the connectivity pattern differs significantly after repetitions, the TTLS melody may not be the sole figure, as the lower voices influence its prominence.

**Figure 1.**
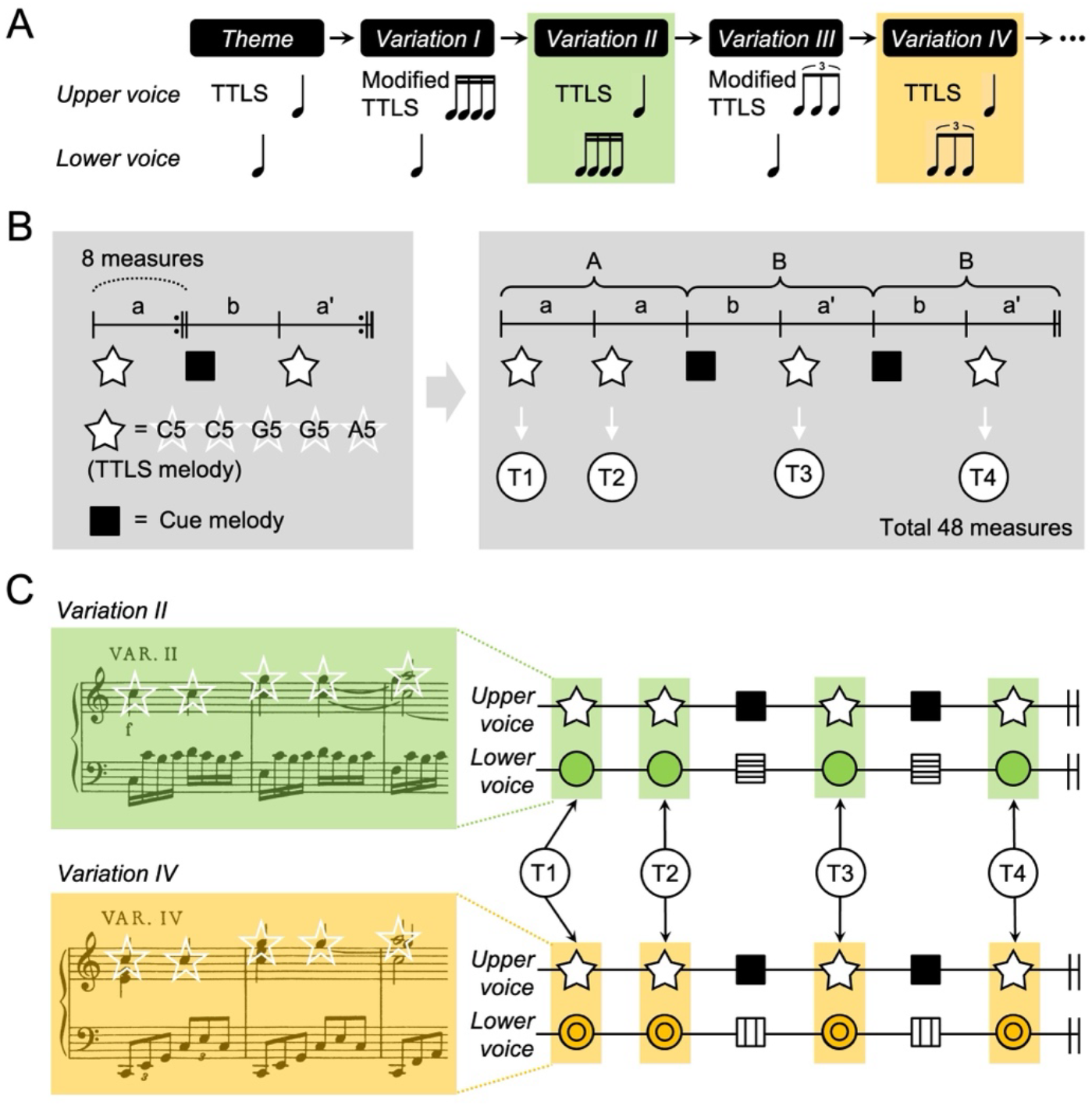
Musical stimuli. **(A)** In Mozart’s 12 Variations, K. 265, the TTLS melody in the theme is modified in *Variations I* and *III* but not in *Variations II* or *IV*. The rhythmic appearance is the same within the pair of *Variations I* and *II* or that of *Variations III* and *IV*. **(B)** The left panel illustrates the structure of each variation involving repeat signs on the score. The right panel depicts the entire structure, comprising 48 measures of A (a + a) + B (b + a’) + B (b + a’), as it is played. White and black squares denote TTLS and cue melodies, respectively, repeated four times per variation. **(C)** The target phrases (T1–T4) are highlighted using green and orange shaded boxes with the “C5-C5-G5-G5-A5” melody marked with white-lined stars. Both *Variations II* and *IV* have two streams of upper and lower voices. The lower voices in the target phrases show melodic and rhythmic variations on the theme, but the upper voice remains consistent. Details about the formation of the adapted and the full scores are shown in Supplementary Figure 3. TTLS, Twinkle Twinkle Little Star; T, target phrase

## Materials and Methods

### Participants

In magnetoencephalography (MEG) recording, participants comprised 25 healthy individuals, all nonmusicians, 15 women and 10 men with a mean age of 26.8 ± 3.4 years old. None had received formal musical training. All participants were right-handed, with a mean Edinburg Handedness coefficient of 95.7 ± 7.1. The study adhered to the principles of the Declaration of Helsinki and received approval from the Institutional Review Board of the Clinical Research Institute at Seoul National University Hospital (IRB No. C-1003-015-311). The research procedures adhered to relevant ethical guidelines and regulations. All participants provided informed, written consent after receiving a clear explanation of the study’s purpose, procedures, potential risks, and benefits.

### Stimuli

Mozart’s K. 265 consists of the theme *‘Ah! Vous dirai-je Maman’* and twelve variations (Supplementary Figure 2). *Variations I–IV* contain rhythmic, melodic, and textural variations on the theme. Relative to the theme, the rhythmic patterns in the upper voices are transformed in *Variations I* and *III* and moved to the lower voices in *Variations II* and *IV*, sharing the TTLS melody (Figure 1A). This study focused on *Variations II* and *IV*, which share the TTLS melody but have differing lower parts, transforming through rhythmic changes to 8^th^ note triplets and 16^th^ notes (semiquavers), respectively. The tonality and harmonic structure remain the same for both variations. Each variation is based on the ternary form of A (a + a) + B (b + a’) + B (b + a’). Phrases, including the “C5-C5-G5-G5-A5” melody, were repeated four times in each variation (Figure 1B). In this study, the term *‘variation’* refers both to the musical form and to individual movements within that form, such as *Variation II* and *Variation IV*.

### Recording

In a magnetically shielded room, the participants listened to Mozart’s K. 265 while watching a silent movie clip (Love Actually, 2003, Universal Pictures, USA) for approximately 5 min. Musical stimuli were generated using STIM^2TM^ (Neuroscan, Charlotte, NC, USA) and presented binaurally at 100 dB through MEG-compatible earphones (Tip-300, Nicolet, Madison, WI, USA). MEG signals were recorded using a 306-channel whole-head MEG system (Elekta Neuromag Vector ViewTM, Helsinki, Finland) with a sampling frequency of 1,000 Hz and a bandpass filter of 0.1–200 Hz. The environmental magnetic noise in raw MEG signals was eliminated using the temporal signal space separation algorithm (Tesche et al., 1995; Taulu and Hari, 2009) implemented in MaxFilter 2.1.13 (Elekta Neuromag Oy, Helsinki, Finland). Electrooculogram, electrocardiogram, and muscle artifacts were also removed using independent component analysis. Participants did not perform behavioral tasks related to attentional shifts between voices during or after MEG recording. They also received no instructions regarding attentional focus during MEG recording.

### Analysis

#### MEG source analysis

The MEG source signals of epochs from −100 ms to 2,100 ms after the onset of each condition for four regional sources of bilateral HGs and IFGs, bandpass filtered at 14–30 Hz, were extracted using BESA 5.1.8.10 (MEGIS Software GmbH, Gräfelfing, Germany) after electrooculograms, electrocardiograms, and muscle artifacts were removed. Standard Talairach coordinates (x, y, and z in millimeters) for bilateral HGs (transverse, BA 41, BA 42) and IFGs (triangular part, BA 45) across participants were adapted from previous research (Kim et al., 2020). The coordinates were as follows: left HG (−53.5, −30.5, and 12.6), right HG (55.4, −30.5, and 12.6), left IFG (−55.5, 11.7, and 20.6), and right IFG (53.5, 12.7, and 20.6).

#### Time window

*Variations II* and *IV* were chosen as the two conditions for estimating connectivity differences, including the identical TTLS melody. Each Time window was 2,100 ms long, incorporating the “C5-C5-G5-G5-A5” melody, as in a previous study (Kim et al., 2020). Time windows of 2,100 ms appeared four times per variation, labeled as T1, T2, T3, and T4 (Figure 1C and Supplementary Figure 2 and 3). The lower voices accompanied by the “C5-C5-G5-G5-A5” melody differed between the two variations.

#### LTDMI analysis

Effective connectivity across target phrases between the two variations was measured using linearized time-delayed mutual information (LTDMI) (Jin et al., 2010; Kim et al., 2021), a measure used in our previous study (Kim et al., 2020). LTDMI estimates the directionality of information transmission between the time series of two regional sources, enabling the observation of interhemispheric and interregional connectivity, which is essential for processing musical elements in bilateral IFGs and HGs. While our primary focus was the regional connection from the left IFG to the right HG, we verified our results for all connections among the bilateral IFGs and HGs. The effective connectivity for the 12 connections between regional sources of the bilateral HGs and IFGs was estimated using MATLAB 7.7.0.471 (Math Works Inc., Natick, MA, USA) (see also Supplementary Table 1 for individual LTDMI values calculated for 12 connections). For each subject, the mean LTDMI for the 2,100-ms epoch was calculated for each of 4 target phrases (T1, T2, T3, and T4) × 2 variations (*Variations II* and *IV)*.

### Statistics

Statistical comparisons of mean LTDMI values for *Variations II* and *IV* were performed using SPSS 21.0 software (IBM, Armonk, NY, USA). For the mean LTDMI values in four target phrases, we conducted the non-parametric Wilcoxon signed-rank test due to the non-Gaussian distribution of LTDMI data. In each case, the significance level (α) for rejecting the null hypothesis (H0, indicating no difference between *Variation II* and *Variation IV* in the mean LTDMI values), was 0.05. In addition, in the nonparametric Spearman correlation test for each pair between the frontotemporal connectivity difference value (Left IFG → Right HG _(*Variation IV*– *Variation II*)_) and the other 11 connectivity difference values, except of Left IFG → Right HG among twelve connections between the bilateral IFGs and HGs, the Type I errors that were caused by multiple comparisons among the 11 connection pairs in the Spearman correlation test were adjusted by the Bonferroni test.

## Results

### LTDMI differences between two variations for four target phrases

The frontotemporal connectivity from the left IFG to the right HG between *Variations II* and *IV* was calculated for four target phrases (T1–T4), each repeated four times per variation (Figure 1). We independently performed a Wilcoxon signed-rank test for the LTDMI values of each target phrase to confirm the changes in the frontotemporal connectivity between the two variations. The difference between the two variations was significant only in T4 among four target phrases, as indicated by the Wilcoxon signed-rank test (Z = −2.112, *P* = 0.035; Figure 2A). In T4, frontotemporal connectivity was enhanced in *Variation IV* compared with *Variation II*. However, significant differences were not observed in T1–T3 (*P* > 0.05 in all cases). In addition, we confirmed that, among 12 connections between the bilateral IFGs and HGs, the only significant result corresponded specially to frontotemporal connectivity from the left IFG to the right HG (Supplementary Table 1). We observed a near-significant effect in temporofrontal connectivity from the right HG to the left IFG, in the opposite direction of frontotemporal connectivity (Right HG → Left IFG, Z = −1.843, *P* = 0.065; Figure 2A and Supplementary Table 1), which was not initially predicted in our hypothesis. The significance level (α) for the null hypothesis was independently tested for each target phrase (T1–T4), since the target phrases of T1–T4 existed in completely different musical contexts within the formal structure of ternary form. Additionally, comparisons between target phrases within a variation were not considered as a hypothesis.

**Figure 2.**
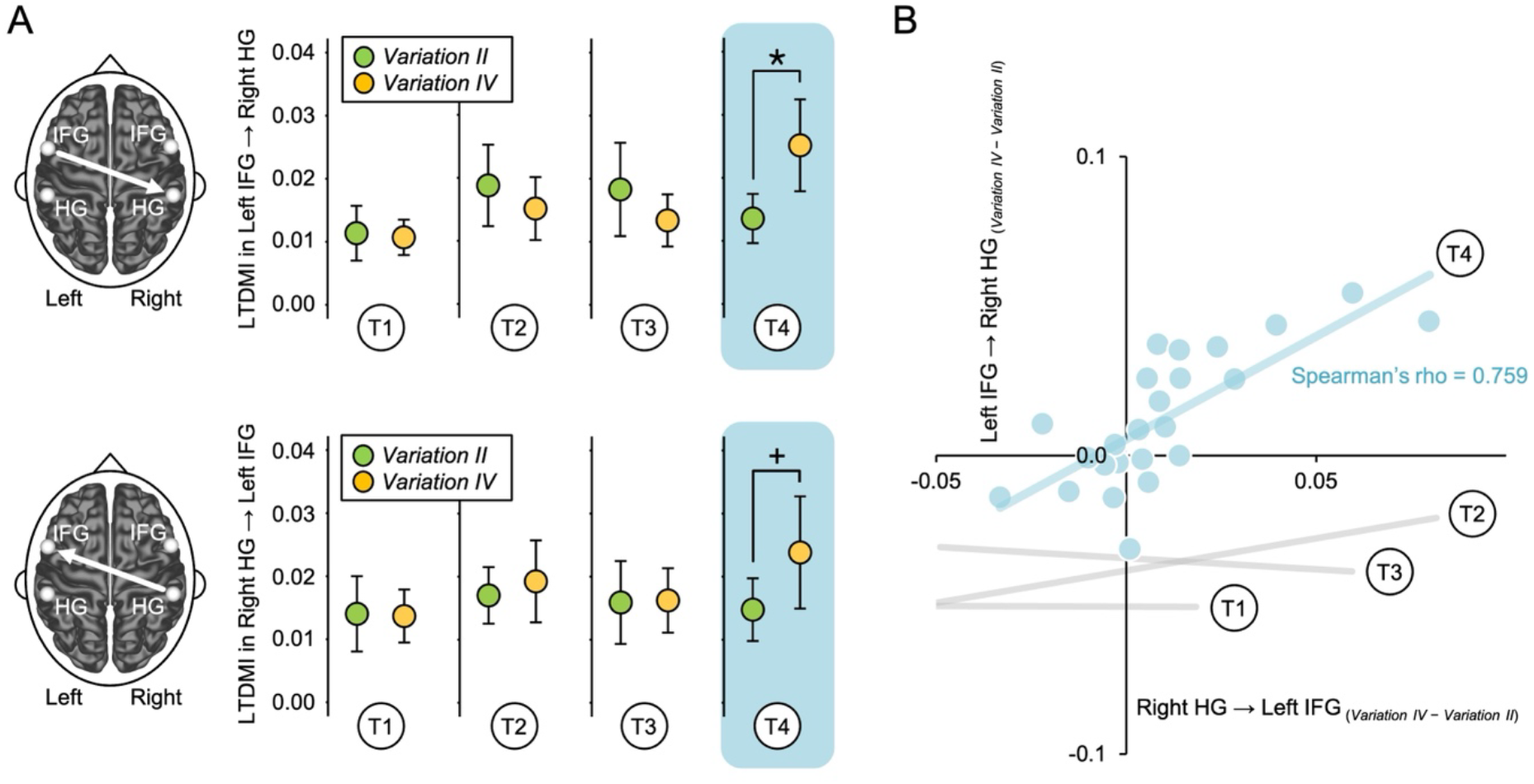
Changes in LTDMI values for four target phrases. **(A)** Frontotemporal connectivity of *Variation IV* was significantly enhanced compared with that of *Variation II* only during T4 (Wilcoxon signed-rank test, Z = −2.112, *P* = 0.035). The temporofrontal connectivity (right HG ⟶ left IFG) of *Variation IV* was also enhanced relative to that of *Variation II* only at T4, although this did not reach the level of statistical significance (Wilcoxon signed-rank test, Z = −1.843, *P* = 0.065). For both frontotemporal and temporofrontal connectivity, there were no significant differences from T1 to T3 (*P* > 0.05 in all cases; Wilcoxon signed-rank test). Error bar denotes the standard error mean. *, *P* < 0.05; +, *P* = 0.65. **(B)** Frontotemporal connectivity (Left IFG → Right HG) was strongly positively correlated with the temporofrontal connectivity (Right HG → Left IFG) only during T4 (Supplementary Table 2). There was a significant correlation between Left IFG → Right HG _(*Variation IV*– *Variation II*)_ and Right HG → Left IFG _(*Variation IV*– *Variation II*)_ at T4 (Spearman correlation, Spearman’s rho = 0.759, Bonferroni-corrected *P* = 0.0001). There were no significant differences from T1 to T3 (*P* > 0.05 in all cases; Spearman correlation; see Supplementary Figure 4). “*Variation IV* – *Variation II*” denotes a difference between *Variation IV* and *Variation II* for the LTDMI value. LTDMI, linearized time delayed mutual information; HG, Heschl’s gyrus; IFG, inferior frontal gyrus; T, target phrase

### Correlation between frontotemporal and temporofrontal connectivity

Correlation analyses were conducted to confirm 1) whether a similar pattern between frontotemporal and temporofrontal connectivity refers to bidirectional information transmission between the left IFG and the right HG and 2) whether a similar pattern is only specialized in the temporofrontal connectivity (Right HG → Left IFG) among 12 connections between the bilateral IFGs and HGs, which are key areas for the music process. To perform this estimation, we first computed the difference values between *Variations II* and *Variation IV* for the LTDMI values in 12 connections between the bilateral IFGs and HGs for all target phrases of T1 to T4. Next, we estimated the correlation between the frontotemporal connectivity difference value (Left IFG → Right HG _(*Variation IV*– *Variation II*)_), with 11 other connectivity difference values. In the Spearman correlation test result, a significant correlation was only observed between Left IFG → Right HG _(*Variation IV*– *Variation II*)_ and Right HG → Left IFG _(*Variation IV*– *Variation II*)_ for T4, among 44 combinations of 11 connections × 4 target phrases (Spearman’s rho = 0.759, Bonferroni-corrected *P* = 0.0001; Figure 2B, Supplementary Figure 4, and Supplementary Table 2). The frontotemporal connectivity (Left IFG → Right HG) was strongly positively correlated with the temporofrontal connectivity (Right HG → Left IFG), reflecting similar information processing in *Variation II* and *IV*.

## Discussion

A difference in frontotemporal connectivity from the left IFG to the right HG between the two variations was only observed in the final target phrase of T4 and not in the preceding three phrases (Figure 2A). As we hypothesized, frontotemporal connectivity showed inconsistency in the figure–ground relationship between the two variations of the TTLS melody in a repeated phrase of T4. This indicates that the perceptual dominance of the TTLS melody in voice perception was weakened. Each variation included cue phrases such as “Up above the world…,” predicting the recurrence of the TTLS melody (Supplementary Figure 2). The two musical structures—variation and ternary forms—establish global and regional contexts, respectively. In each variation, T4 is introduced after the second iteration of the cue phrase within the regional ternary context, facilitating anticipation of melodic recurrence. Moreover, at the global level of the variation form, the same structure involving T4 is repeated in *Variation II* and *Variation IV*, further enhancing anticipatory processing. The training of the repeated upper voice and its perceptual prominence might have facilitated participants’ recognition of the lower voice (Taher et al., 2016). The LTDMI value was higher in *Variation IV* than in *Variation II* during T4. We interpret that the connectivity reduction in *Variation II* for T4 is attributable to the properties of its lower voice, which differed from them in *Variation IV* (Figure 3). Our findings show that participants did not solely focus on the TTLS melody at T4 but could also detect other sounds.

**Figure 3.**
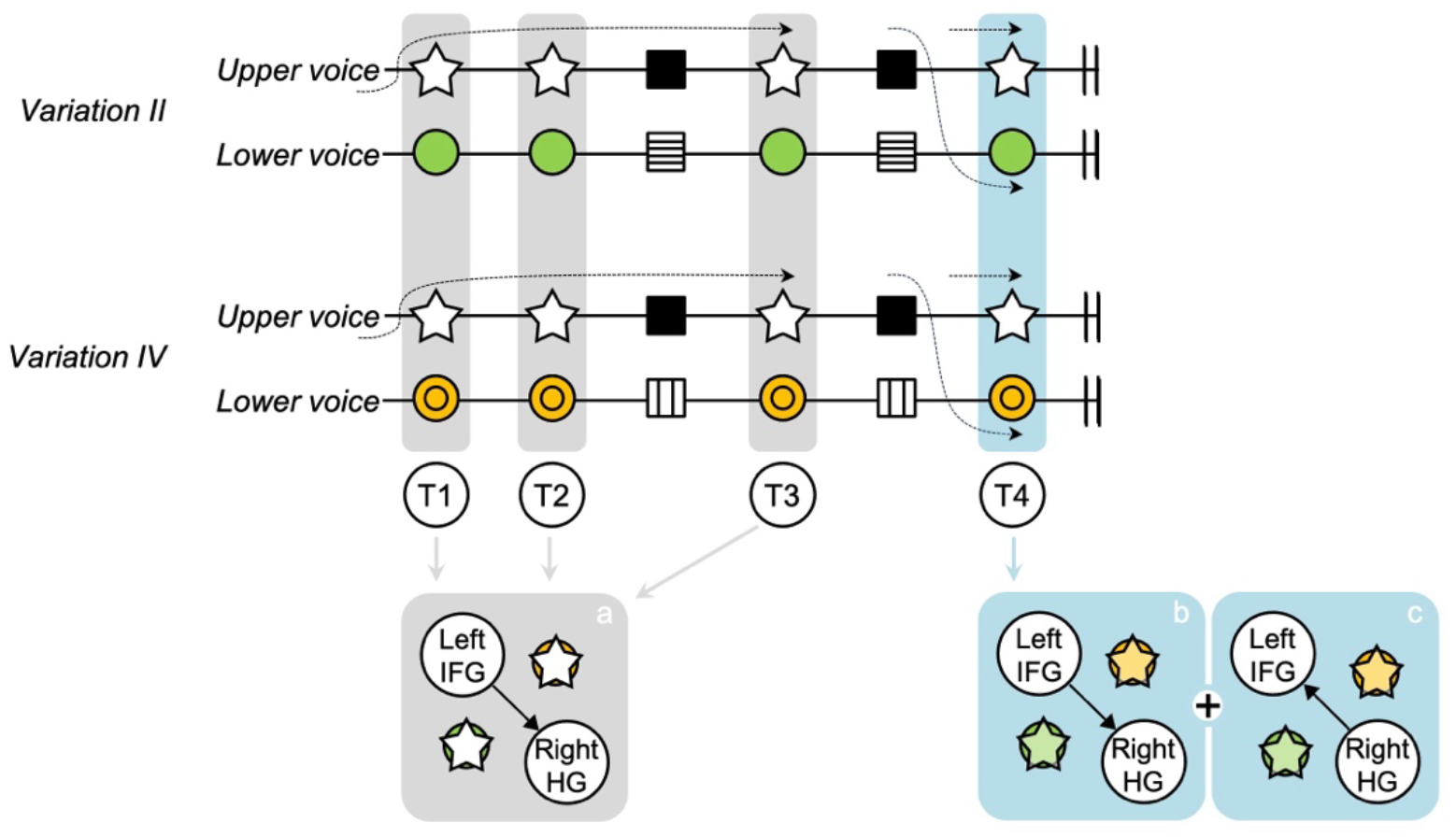
Figure–ground relationship between voices according to frontotemporal and temporofrontal connectivity changes. The figure shows how connectivity between the left IFG and the right HG changes from T1 to T4 when the same passages are repeated in each variation. Frontotemporal connectivity from the left IFG to the right HG, the TTLS connectivity, is consistent from T1 to T3, focusing on the figure of the upper voice. However, at T4, a different frontotemporal connectivity pattern is exhibited between *Variations II* and *IV*. This indicates that frontotemporal connectivity may no longer be related to the TTLS melody but rather to the lower voice, reflecting a shift away from TTLS-specific connectivity. Lower voices, masked by the perceptual dominance of the TTLS melody until T3, may have become audible alongside the TTLS melody at T4. However, because these results do not demonstrate whether figure and ground are perceptually separated or integrated, predictable bidirectional processes are represented using overlapping circles and stars. HG, Heschl’s gyrus; IFG, inferior frontal gyrus; T, target phrase; TTLS, Twinkle Twinkle Little Star

In our previous study (Kim et al., 2020), unidirectional transmission of information from the left IFG to the right HG, showing frontotemporal connectivity, was associated with the recognition of the TTLS melody in the heading of each variation. However, the involvement of temporofrontal connectivity, which was in the opposite direction to the frontotemporal connectivity, was observed in repeated phrases in musical context evoked (Figure 2B). The temporofrontal connectivity was strongly correlated with the frontotemporal connectivity in T4 (Figure 2B and Supplementary Table 2). The top–down processing of a familiar melody can significantly influence the figure–ground relationship (Strüber and Stadler, 1999; Nelson and Palmer, 2007). Considering that the roles of frontotemporal and temporofrontal connectivity are linked to the TTLS melody-based and sensory-based processes, respectively, this heightened connectivity could indicate a dual process: extracting the novel lower voice in the target phrase, including the familiar TTLS melody, and dissecting the components within the novel lower voices. The temporofrontal connectivity supports the frontotemporal connectivity. Thus, bidirectional connectivity of the frontotemporal and temporofrontal pathways between the left IFG and the right HG is possibly modulated by both a top–down process based on knowledge of the TTLS melody and a bottom–up process based on new information on the voices accumulated while sequentially listening to target and cue phrases in each variation (Alho et al., 2015; Dzafic et al., 2021).

Numerous researches on the auditory figure–ground relationship have conducted auditory scene analysis (Bregman, 1994) and grouping (Wagemans et al., 2012) using tasks that discriminate a sound pattern as a figure from a ground of tone and chord sequences with irregularities in spectral and temporal properties (Teki et al., 2011; O’Sullivan et al., 2015; Toth et al., 2016). Functional magnetic resonance imaging and electroencephalography studies (Teki et al., 2011; O’Sullivan et al., 2015; Toth et al., 2016) have reported that regions involved in discriminating this figure–ground perception comprise the primary auditory area, superior temporal sulcus, superior temporal gyrus, intraparietal sulcus, medial/superior frontal gyrus, and cingulate cortex. The processing of multiple voices in this study involved the IFG and the HG, both exhibiting information transmission. The left IFG is crucial in processing familiarity (Plailly et al., 2007) and in music-syntactic processing, indicating implicit learning (Sammler et al., 2011) and contributing to memory retrieval (Platel et al., 2003; Watanabe et al., 2008). Moreover, the left IFG is involved in differentiating between melody and accompaniment (Spada et al., 2014). The left IFG is highly activated during the recognition of pattern deviations in melody (Habermeyer et al., 2009), musical novelty in a context (Tillmann et al., 2003), and conscious experience (Weilnhammer et al., 2021). The left IFG is also associated with sematic and syntactic processing (Zhu et al., 2022), and sentence comprehension (van der Burght et al., 2019). In contrast, the right HG predominantly processes tone deviance (Sabri et al., 2006; Nan and Friederici, 2013) and the segregation of auditory streams (Snyder et al., 2006). The right auditory cortex is the dominant site for music processing (Perani et al., 2010), auditory stream segregation (Snyder et al., 2006), and spectral pitch (Schneider et al., 2005).

The IFG and HG are pivotal areas for music perception, and their connectivity is discussed in relation to syntax processes (Papoutsi et al., 2011; Kim et al., 2019; Kim et al., 2021), categorization (Roswandowitz et al., 2021), and the working memory of the melody process (Burunat et al., 2014). The temporofrontal network is engaged in the categorization of vocal signals (Roswandowitz et al., 2021), while frontotemporal connection is involved in both top-down and bottom-up processes in sensory learning (Dzafic et al., 2021). Directional information flows within frontotemporal connectivity may therefore explain how the left IFG and the right HG collaborate in processing target phrases. The involvement of the left IFG and right HG may reflect the entire process of naturally grasping, comparing, and understanding the voices in target phrases within a particular context rather than simply perceiving them as sounds. Accordingly, enhanced frontotemporal and temporofrontal connectivity may reflect the integrated processes by which the brain recognizes the TTLS melody based on memory, segregates the melody, and detects differences in the lower voices relative to the prior context.

Previous studies have selectively manipulated stimuli or directed participants’ attention to specific auditory streams (Uhlig et al., 2013; Ragert et al., 2014; Spada et al., 2014; Strait et al., 2015; Hausfeld et al., 2018; Puschmann et al., 2019; Barrett et al., 2021). Attention has been shown to be critical for figure–ground perception (Poort et al., 2012). Therefore, research on such perception uses artificially composed stimuli to direct participants’ attention. During the MEG experiment in this study, all participants passively listened to the naturalistic music of Mozart’s 12 Variations, K. 265, without any instructions regarding focusing their attention on a specific voice or melody. Thus, whether both figure and ground were processed attentively or pre-attentively remains unclear, given the absence of intentional attention control. Although listeners may focus on a particular voice while listening to music, the changing flow of music can encompass brief perceptible moments in which the figure– ground relationship continuously shifts without listeners consciously realizing it. Indeed, music listeners can automatically process information such as syntactic errors and tone deviations without intentional attention (Maess et al., 2001; Naatanen et al., 2007). We interpreted that participants could attentively or pre-attentively detect sonic changes in the voices at that moment, although our data do not confirm that non-musicians could perceptually segregate the streams or identify which voice evoked the sonic differences. (Figure 3). Our results successfully captured the moment when the figure–ground relationship between the upper and lower voices changed, as evidenced by the difference in frontotemporal connectivity for repeated phrases in the two variations and the correlation between frontotemporal and temporofrontal connectivity.

Familiarity is critical for explaining the figure–ground experiment (Palmer, 1999; Hulleman and Humphreys, 2004; Nelson and Palmer, 2007). The target phrases in our stimuli involved the TTLS song, which has been used in studies related to the perception of familiar melodies (Trehub et al., 1985; Upitis, 1990; Besson et al., 1994; Creel, 2019). Familiarity would be naturally implied in the theme and all of its variations, considering that Mozart’s 12 Variations, K. 265, is based on the TTLS melody. Logically, the familiarity implied in the TTLS melody might influence participants’ figure–ground perception. However, in our previous study focusing on the TTLS melody (Kim et al., 2020), we could not directly prove the effect of familiarity on frontotemporal connectivity as the connectivity changed irrespective of the presence or absence of the TTLS melody. In our present study, the same TTLS melody appeared repeatedly in *Variations II* and *IV*. The effects of familiarity are consistent in both *Variations II* and *IV*. Thus, the TTLS melody was used to assess changes in the figure–ground relationship.

Naturalistic stimuli have been used to examine various topics (Saarimäki, 2021; Izen et al., 2023; Tervaniemi, 2023). In studies on the concepts of emotion (Singer et al., 2016; Putkinen et al., 2021), melodic expectation (Kern et al., 2022), temporal aspects of rhythm and beat (Sturm et al., 2015; Weineck et al., 2022), and familiarity (Leaver et al., 2009), multiple naturalistic pieces have been used as musical stimuli. Some studies using naturalistic stimuli have examined their hypotheses on topics such as motif, musical features, timbre, and depression, based on a single piece (Alluri et al., 2012; Cong et al., 2013; Burunat et al., 2016; Liu et al., 2020). Our hypothesis was also created for the melody of TTLS and the figure–ground relationship of voices using Mozart’s 12 Variations, K. 265. In the fields of audiation (Uhlig et al., 2013; Ragert et al., 2014; Hausfeld et al., 2018; Barrett et al., 2021) and vision (Peterson et al., 1991; Lamme, 1995; Super et al., 2003; Zhang and Von Der Heydt, 2010; Von der Heydt, 2015), changes in the figure–ground relationship between the elements of an object can be simply observed and explained by comparing the related objects. However, naturalistic music has its own narrative, which can be described by its structure.

The connectivity reflects complicated processes for the target phrases of 2.1 s without the context and for each 2.1-s-long target phrases in the theme and variations, leading up to the target phrase. This approach, however, may constitute a critical shortcoming of our study, compared with measurements using artificially composed stimuli. Furthermore, no verbal reports or other measures were obtained during or after MEG recording to determine what participants perceived as the ‘figure’ in the music at each moment, including critical time windows. As a result, connectivity changes were measured under naturalistic listening conditions in which participants passively listened to music. These limitations can be addressed in future studies using novel paradigms that incorporate detailed behavioral responses and larger sample sizes. Nevertheless, our results reflected the humans’ ubiquitous experiences with individual participants.

In addition to the use of naturalistic music, our study had other limitations. While non-musicians can perceive the figure separately from the ground (Toth et al., 2016), they tend to focus more on the upper voice than on a lower one (Sloboda and Edworthy, 1981). In contrast, musicians are more sensitive to voice perception and are better able to distinguish voices (Fujioka et al., 2005; Strait et al., 2015). Music training significantly influences selective attention (Puschmann et al., 2019). We did not recruit musicians as participants to examine our hypothesis in terms of basic musical ability. Recruiting non-musicians might have impacted our results. The temporofrontal connectivity may show a statistically significant distinction (*P* < 0.05) since the recognition of different elements in the lower voices (via temporofrontal connectivity) involves a more complex cognitive process. Therefore, future studies should verify these results with musicians. Furthermore, this study concentrated on frontal and temporal regions. Our findings should be verified at the whole-brain level. In the current experimental paradigm, we did not consider an additional test individual preference of subjects. The preference is also influential of music listening, and which should be addressed importantly in further studies with a novel experimental paradigm. The MEG recording and analysis approach used in this study may be replicable using other modalities, such as EEG. Despite these limitations, the use of Mozart’s 12 Variations, K. 265, was invaluable in understanding the fundamental neural processes associated with processing real music featuring multiple voices and elucidating the human experience of music. Our findings elucidated how the brain dissects voices from the multidimensional structures of music and reconstructs the figure–ground relationship between voices.

## Acknowledgment

We sincerely appreciate Ji Hyang Nam for her technical support in MEG data acquisition.

## Author contributions

CHK: Conceptualization, Formal analysis, Funding acquisition, Supervision, Methodology, Visualization, Writing – original draft, Writing – review & editing; JES: Methodology, Writing – review & editing; JS: Investigation, Methodology, Writing – review & editing; CCK: Funding acquisition, Supervision, Writing – review & editing

