## Supplementary figures and tables for "Figure–ground relationship of voices in musical structure modulates reciprocal frontotemporal connectivity"

### Supplementary materials

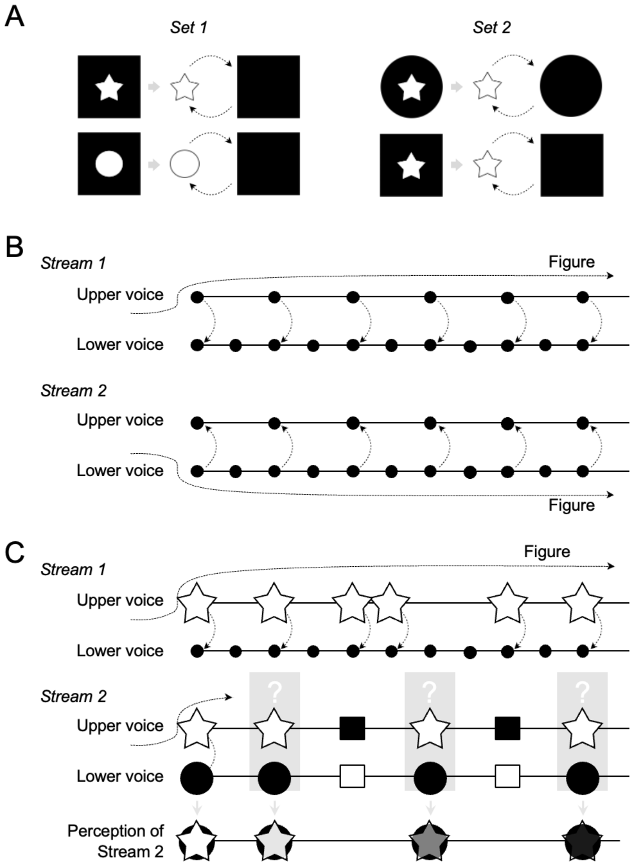


**Supplementary Figure 1. Figure–ground perception in visual and auditory stimuli. (A)** Sets 1 and 2 demonstrate examples of figure–ground perception in visual stimuli. In Set 1, both objects are presented against a shared black square background. Here, the black squares denote figures and the white star and circle represent the ground. In Set 2, the white stars denote figure and the black circle and square represent the ground. This relationship can also be reversed. The primary distinction between the two objects lies in the differing shapes of the black circle and square. **(B)** In polyphonic music, as illustrated in Stream 1, the lower voice within a phrase functions as the ground, while the listener's attention is drawn to the upper voice, perceived as the figure. In Stream 2, these roles are reversed, with the lower voice becoming the figure. This model is adapted from Bigand et al. (2000). Even if the figure–ground relationship shifts, the presence of the ground does not indicate that the figure is the only perceived sound. As indicated by the dotted arrows between the voices, a part of the ground may still be heard, albeit faintly. **(C)** When a familiar melody (white stars) appears within the musical piece, it is easily recognized as the figure (Stream 1). However, with repeated exposure to music with recurring phrases (black squares), the perceptual prominence of the figure might be diminished (Stream 2). Consequently, in musical structure, the figure–ground relationship centered on the upper voice may begin to shift, potentially leading to a natural collapse of the distinction between the figure and the ground.

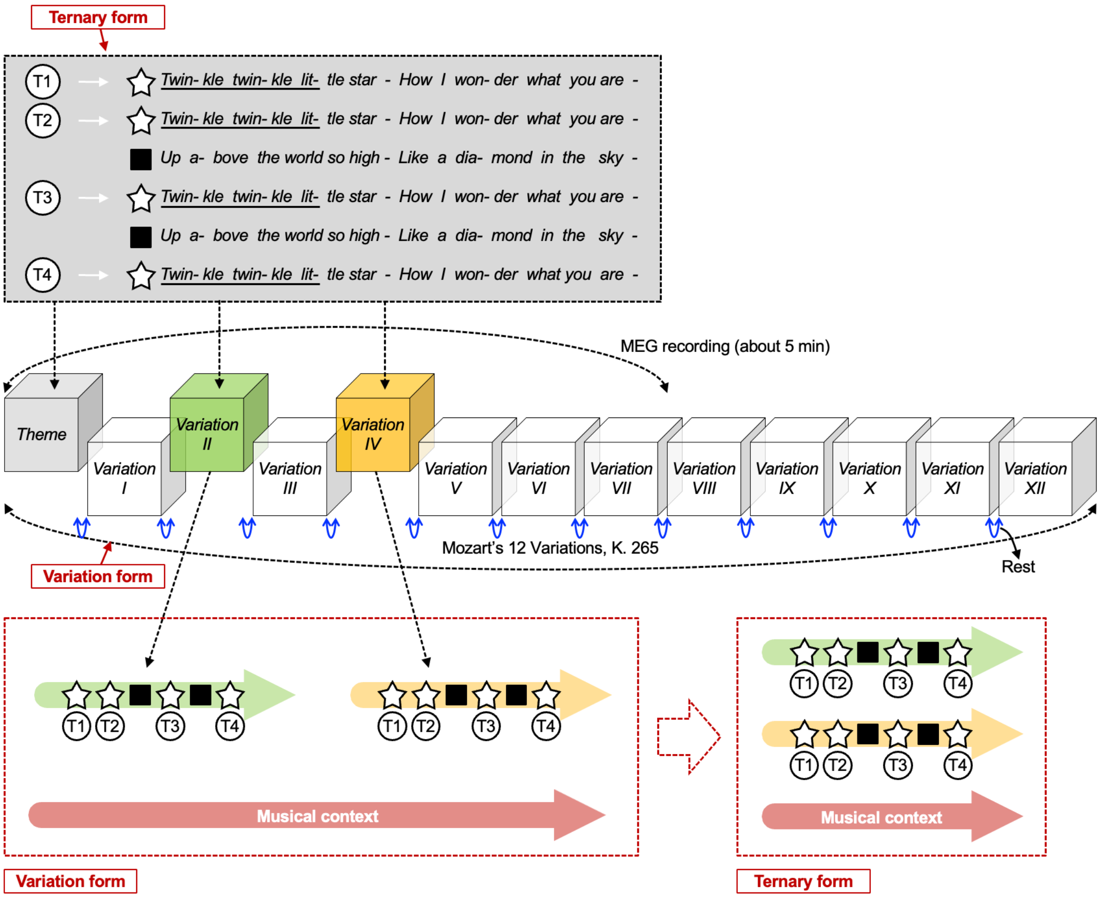


**Supplementary Figure 2. Musical stimuli and structure.** Mozart's K. 265 consists of a theme and twelve variations based on variation form. During MEG recording, participants listened to approximately 5 minutes of the piece, ending near the conclusion of *Variation VII*. Among the twelve variations, we selected *Variation II* and *Variation IV* for analysis because they share the same musical structure: a ternary form, A (a + a) + B (b + a') + B (b + a'), and both retain the original melody of *"Twinkle, Twinkle, Little Star."* Within this piece—particularly from the *Theme* through *Variation IV*—two types of musical structures are present: variation form at the level of the entire piece and ternary form at the level of individual movements (variations). The time windows containing the melody "C5–C5–G5–G5–A5" are repeated four times (T1, T2, T3, and T4) in each variation (Figure 1B). The lyrics of *Twinkle, Twinkle, Little Star* can be adapted to this ternary form. The repeating phrase "Up above the world…" (black squares) facilitates prediction of the reappearance of T3 and T4 (white stars) as cue melodies. The cue melody plays a stronger role in T4, where it reappears, than in T3. Musical context is present in both variation and ternary forms. The theme and twelve variations are separated by rests across the variation form. In the context of the entire piece, effects related to repetition are expected to be more pronounced in *Variation IV* than in *Variation II* due to repeated structural similarity. Within the ternary form, the regional structure from T1 to T4 is synchronized and parallel between *Variation II* and *Variation IV*. Our hypotheses and statistical analyses focused on this synchronized regional context, while also potentially reflecting global variation-level context. In this study, the term *"variation"* has two meanings: (1) the musical form and (2) individual movements such as *Variation II* and *Variation IV*. See Supplementary Figure 3 for the musical score from the *Theme* through *Variation IV*.

### Zwölf Variationen in C

über das französische Lied «Ah, vous dirai-je Maman»

KV 265 (300^e)

Endstanden wahrscheinlich Paris, 1778

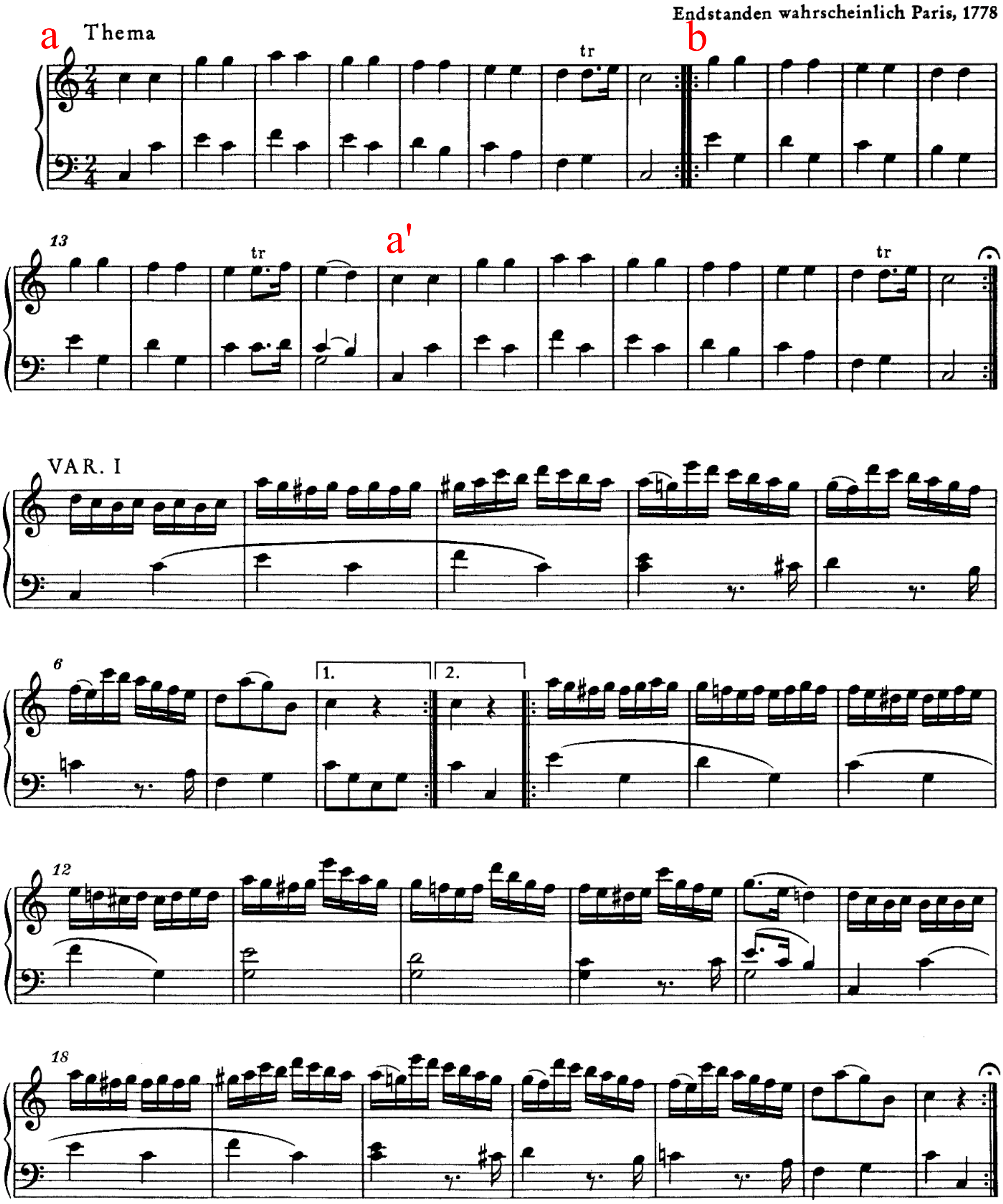

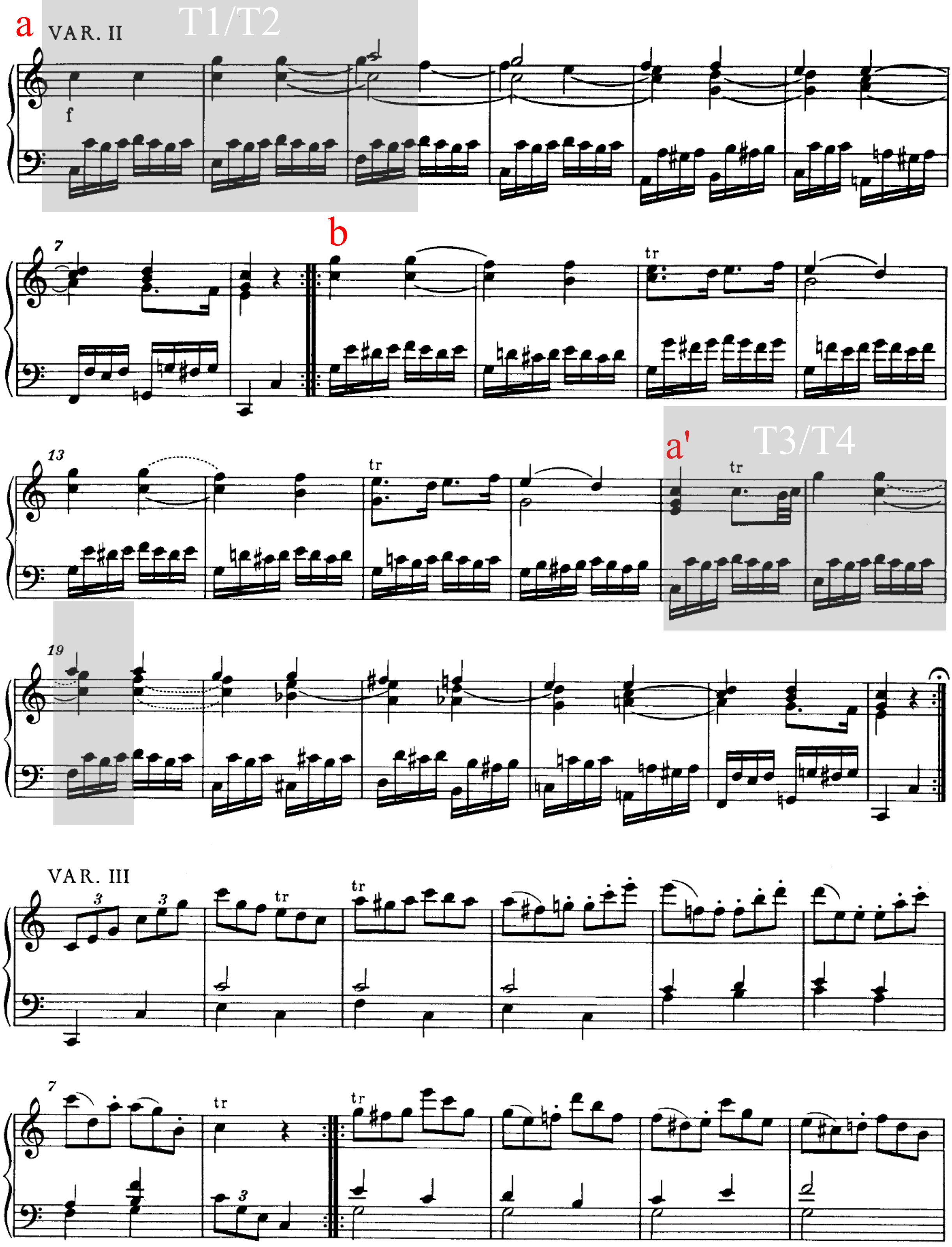
a
VAR. II
T1/T2
f
7
b
tr
13
a'
tr
T3/T4
19
VAR. III
3
7

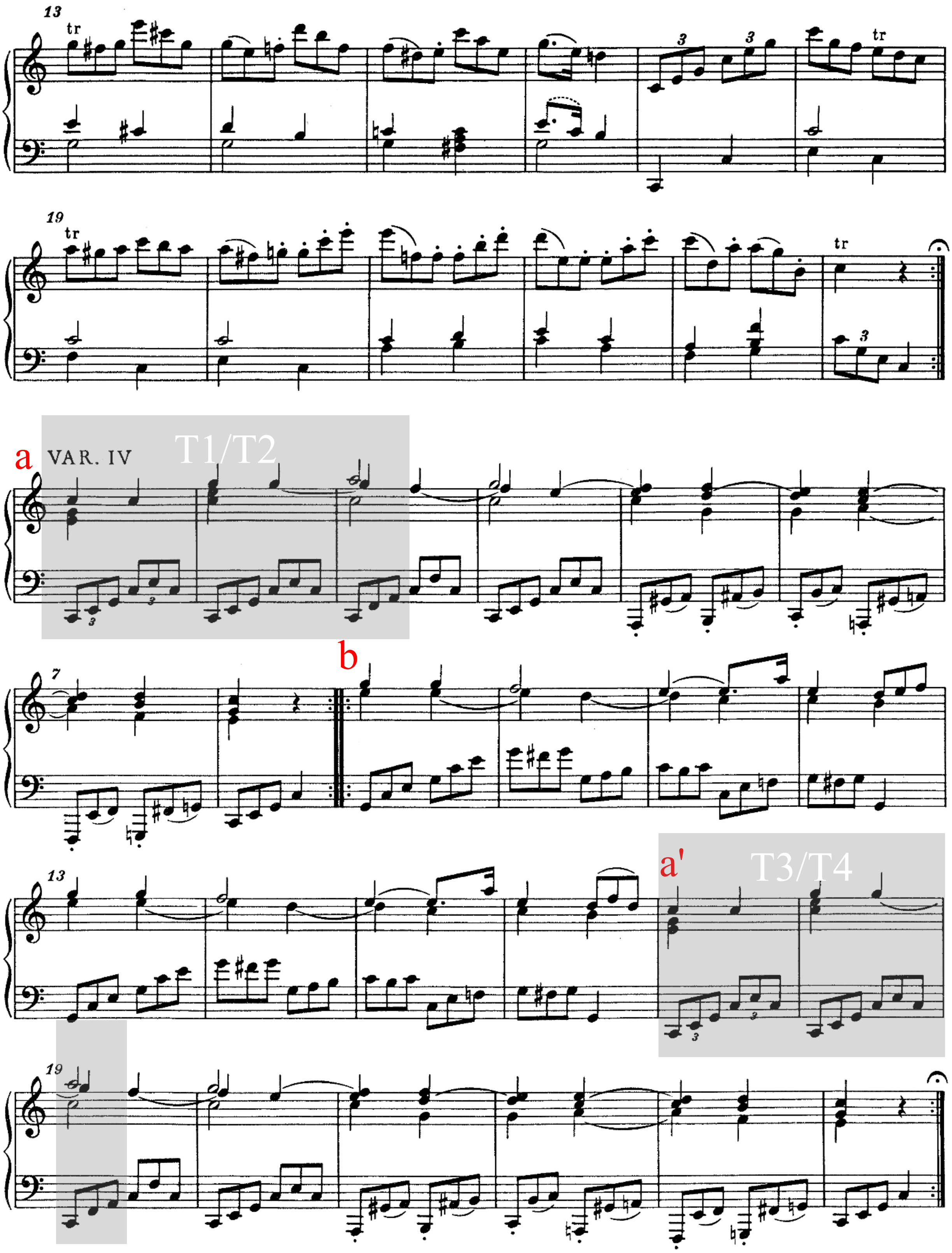




**Supplementary Figure 3. Musical stimulus.** Musical score of the theme to *Variation IV* in Mozart's 12 Variations on "Ah, vous dirai-je maman" K. 265 (Barenreiter edition) was adapted from *NMA Online: Neue Mozart-Ausgabe: Digitized Version* (https://dme.mozarteum.at/DME/nma/nmapub_srch.php?l=2). Each movement is based on the ternary form of A (a + a) + B (b + a') + B (b + a'). The time windows of T1, T2, T3, and T4 in *Variations II* and *IV* are marked by gray shaded boxes.

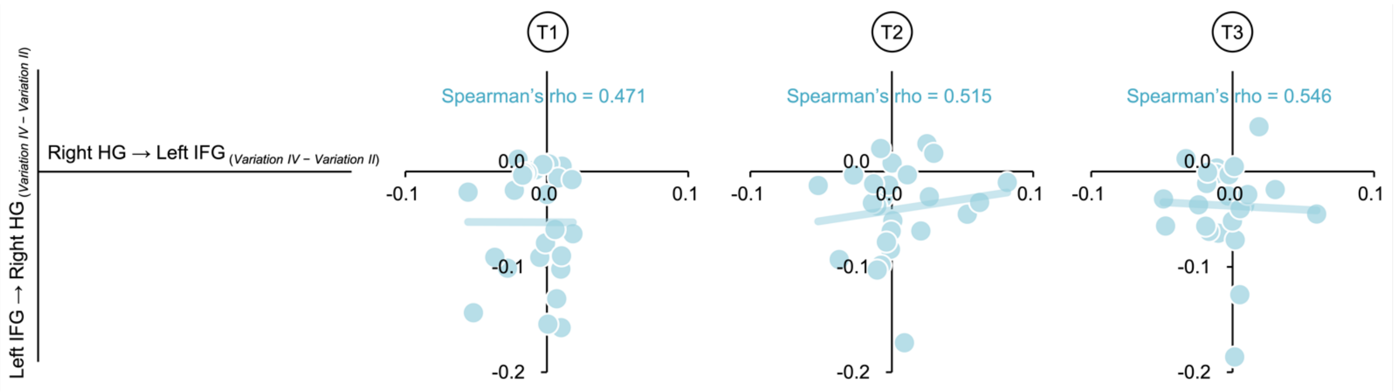


**Supplementary Figure 4. Correlation between frontotemporal and temporofrontal connectivity from T1 to T3.** See Supplementary Table 2 for the detail for the statistics.

**Supplementary Table 1. Difference in mean LTDMI values between *Variations II* and *IV* across four target phrases for 12 connections between the bilateral HGs and IFGs.** We tested for 12 connections between the bilateral HGs and IFGs to confirm whether our hypothesis was valid, even though our hypothesis was limited to one connection among the 12 connections. Significance was observed in a single connection from the left IFG to the right HG in T4 among 48 combinations of 12 connections and four target phrases. A tendency toward significance was observed in a connection from the right HG to the left IFG. *, $P < 0.05$.

| | **T1** | | **T2** | | **T3** | | **T4** | |
|---|---|---|---|---|---|---|---|---|
| | **Z** | **P** | **Z** | **P** | **Z** | **P** | **Z** | **P** |
| **Left HG → Right HG** | -0.296 | 0.767 | -0.578 | 0.563 | -0.632 | 0.527 | -0.632 | 0.527 |
| **Left HG → Left IFG** | -0.498 | 0.619 | -1.278 | 0.201 | -0.874 | 0.382 | -0.901 | 0.367 |
| **Left HG → Right IFG** | -0.794 | 0.427 | -0.794 | 0.427 | -1.682 | 0.093 | -0.336 | 0.737 |
| **Right HG → Left HG** | -1.278 | 0.201 | -0.363 | 0.716 | -1.332 | 0.183 | -0.283 | 0.778 |
| **Right HG → Left IFG** | -0.296 | 0.767 | -0.148 | 0.882 | -0.256 | 0.798 | ***-1.843*** | ***0.065*** |
| **Right HG → Right IFG** | -0.659 | 0.510 | -1.574 | 0.115 | -1.063 | 0.288 | -0.363 | 0.716 |
| **Left IFG → Left HG** | -0.013 | 0.989 | -0.982 | 0.326 | -1.036 | 0.300 | -1.574 | 0.115 |
| **Left IFG → Right HG** | -0.148 | 0.882 | -0.767 | 0.443 | -0.767 | 0.443 | ***-2.112*** | ***0.035 **** |
| **Left IFG → Right IFG** | -1.682 | 0.093 | -0.928 | 0.353 | -0.229 | 0.819 | -0.578 | 0.563 |
| **Right IFG → Left HG** | -1.009 | 0.313 | -0.632 | 0.527 | -0.013 | 0.989 | -0.067 | 0.946 |
| **Right IFG → Right HG** | -0.027 | 0.979 | -1.440 | 0.150 | -0.417 | 0.677 | -1.090 | 0.276 |
| **Right IFG → Left IFG** | -1.413 | 0.158 | -0.928 | 0.353 | -1.762 | 0.078 | -0.484 | 0.628 |

**Supplementary Table 2. Correlation between frontotemporal connectivity and other connectivity for the difference values of *Variation II* and *IV*.** We first computed the differences in value between *Variations II* and *IV* (*Variation IV– Variation II*) for the LTDMI values in 12 connections for the bilateral IFGs and HGs for all target phrases of T1 to T4. Next, we estimated the correlations between frontotemporal connectivity difference value (Left IFG → Right HG $_{(Variation\ IV-\ Variation\ II)}$) and other 11 values for the connectivity difference. In the Spearman correlation test result, the significant positive correlation was only observed between Left IFG → Right HG $_{(Variation\ IV-\ Variation\ II)}$ and Right HG → Left IFG $_{(Variation\ IV-\ Variation\ II)}$ for T4. Type I errors caused by multiple comparisons between the 11 connection pairs for each target phrase in the Spearman correlation test were adjusted with the Bonferroni test.

| | A | B | Correlation between A and B | | |
|---|---|---|---|---|---|
| **Target phrase** | **Difference value of "*Variation IV – Variation II*"** | **Difference value of "*Variation IV – Variation II*"** | **Spearman's rho** | ***P*** | **Bonferroni-corrected *P*** |
| ***T1*** | Left IFG → Right HG | Right HG → Left IFG | 0.471 | 0.018 | 0.193 |
| | | Left HG → Right STG | 0.000 | 0.999 | 10.984 |
| | | Left HG → Left IFG | 0.147 | 0.483 | 5.318 |
| | | Left HG → Right IFG | -0.200 | 0.338 | 3.716 |
| | | Right HG → Left HG | 0.157 | 0.453 | 4.978 |
| | | Right HG → Right IFG | -0.113 | 0.590 | 6.495 |
| | | Left IFG → Left HG | 0.142 | 0.500 | 5.497 |
| | | Left IFG → Right IFG | 0.160 | 0.444 | 4.880 |
| | | Right IFG → Left HG | -0.267 | 0.197 | 2.168 |
| | | Right IFG → Right HG | 0.068 | 0.745 | 8.196 |
| | | Right IFG → Left IFG | 0.077 | 0.715 | 7.862 |
| ***T2*** | Left IFG → Right HG | Right HG → Left IFG | 0.515 | 0.008 | 0.092 |
| | | Left HG → Right STG | 0.293 | 0.155 | 1.706 |
| | | Left HG → Left IFG | 0.145 | 0.490 | 5.394 |
| | | Left HG → Right IFG | -0.001 | 0.996 | 10.952 |
| | | Right HG → Left HG | 0.402 | 0.046 | 0.510 |
| | | Right HG → Right IFG | -0.216 | 0.299 | 3.293 |
| | | Left IFG → Left HG | 0.005 | 0.980 | 10.776 |
| | | Left IFG → Right IFG | 0.074 | 0.724 | 7.967 |
| | | Right IFG → Left HG | -0.116 | 0.582 | 6.397 |
| | | Right IFG → Right HG | -0.052 | 0.807 | 8.874 |
| | | Right IFG → Left IFG | -0.185 | 0.375 | 4.124 |
| ***T3*** | Left IFG → Right HG | Right HG → Left IFG | 0.546 | 0.005 | 0.053 |
| | | Left HG → Right STG | 0.379 | 0.061 | 0.676 |
| | | Left HG → Left IFG | -0.457 | 0.022 | 0.239 |
| | | Left HG → Right IFG | 0.112 | 0.595 | 6.550 |
| | | Right HG → Left HG | 0.222 | 0.286 | 3.149 |
| | | Right HG → Right IFG | 0.163 | 0.437 | 4.808 |
| | | Left IFG → Left HG | -0.172 | 0.411 | 4.522 |
| | | Left IFG → Right IFG | 0.435 | 0.030 | 0.329 |
| | | Right IFG → Left HG | 0.220 | 0.290 | 3.187 |
| | | Right IFG → Right HG | 0.189 | 0.365 | 4.014 |
| | | Right IFG → Left IFG | 0.546 | 0.005 | 0.052 |
| ***T4*** | **Left IFG → Right HG** | **Right HG → Left IFG** | ***0.759*** | ***0.00001*** | ***0.0001 ****** |
| | | Left HG → Right STG | 0.037 | 0.861 | 9.470 |
| | | Left HG → Left IFG | -0.255 | 0.218 | 2.396 |
| | | Left HG → Right IFG | -0.397 | 0.049 | 0.544 |

| | | | |
|---|---|---|---|
| Right HG → Left HG | 0.013 | 0.951 | 10.456 |
| Right HG → Right IFG | 0.188 | 0.369 | 4.058 |
| Left IFG → Left HG | -0.304 | 0.140 | 1.538 |
| Left IFG → Right IFG | 0.075 | 0.722 | 7.937 |
| Right IFG → Left HG | -0.022 | 0.919 | 10.105 |
| Right IFG → Right HG | 0.155 | 0.460 | 5.065 |
| Right IFG → Left IFG | 0.224 | 0.282 | 3.103 |
